# Distinct photooxidation-induced cell death pathways lead to selective killing of human breast cancer cells

**DOI:** 10.1101/2020.06.25.170860

**Authors:** Ancély F. dos Santos, Alex Inague, Gabriel S. Arini, Letícia F. Terra, Rosangela A.M. Wailemann, André C. Pimentel, Marcos Y. Yoshinaga, Ricardo R. Silva, Divinomar Severino, Daria Raquel Q. de Almeida, Vinícius M. Gomes, Alexandre Bruni-Cardoso, Walter R. Terra, Sayuri Miyamoto, Maurício S. Baptista, Leticia Labriola

## Abstract

Lack of effective treatments for aggressive breast cancer is still a major global health problem. We previously reported that Photodynamic Therapy using Methylene Blue as photosensitizer (MB-PDT) massively kills metastatic human breast cancer, marginally affecting healthy cells. In this study we aimed to unveil the molecular mechanisms behind MB-PDT effectiveness. Through lipidomic and biochemical approaches we demonstrated that MB-PDT efficiency and specificity relies on polyunsaturated fatty acids-enriched membranes and on the better capacity to deal with photooxidative damage displayed by non-tumorigenic cells. We found out that, in tumorigenic cells, lysosome membrane permeabilization is accompanied by ferroptosis and/or necroptosis. Our results broadened the understanding of MB-PDT-induced photooxidation mechanisms and specificity in breast cancer cells. Therefore, we demonstrated that efficient approaches could be designed on the basis of lipid composition and metabolic features for hard-to-treat cancers. The results further reinforce MB-PDT as a therapeutic strategy for highly aggressive human breast cancer cells.

## Introduction

Breast cancer is the most frequent malignancy in women worldwide^1,2^. In its advanced stages, when distant organ metastases occur, it is considered incurable with the currently available therapies^2^. The reason being that metastatic lesions are usually multiple, molecular and cellular heterogenous, and resistant to conventional treatments^3^. Thus, effective and safe therapies for this stage of the disease are still needed.

Photodynamic therapy (PDT) has been the focus of several cancer centers as it might represent an important advancement in treatment due to its high but also controlled cytotoxic effect^4^. Additionally, the enhanced antitumor effects combining PDT and chemotherapies have already been demonstrated in preclinical studies on breast cancer^3^. PDT consists in the uptake of a photosensitizer (PS) molecule which, upon excitation by light in a determined wavelength, reacts with oxygen and generates oxidant species (radicals, singlet oxygen, triplet species) in target tissues, leading to photooxidative stress (PhOxS)^5,6^, which results in photodamage of membranes and organelles^7,8^. The extent of the damage, and the cell death mechanisms involved, are dependent on the PS type, concentration, subcellular localization, the amount of energy and fluence rate applied as well as on the intrinsic characteristics of each tumor type^9–12^. The bottleneck of PDT is that little is known about the complex molecular mechanisms behind its cytotoxicity and even less about the factors that could improve its specificity against aggressive cancer cells. In order to address these underpinnings, our group has been studying PDT using methylene blue as photosensitizer (MB-PDT) in human breast cell (BC) models.

In previous studies we have already reported that there were differences in MB-PDT sensitivity regarding MB concentration, time to achieve maximal cell death and the effect of fluence rate^9,13^. Moreover, our results have shown that non-tumorigenic breast cells are more resistant to MB-PDT, whereas the very aggressive triple negative breast cancer cells (TNBC) displayed the highest susceptibility^13^. However, the mechanisms behind these effects are still not well understood. In the present study, we set out to unveil the molecular mechanisms triggered by this PhOxS therapy that are responsible for its selectivity in the elimination of human breast cancer cells. For this purpose we performed a comprehensive and comparative lipidomic profiling of two breast cancer cell types (MDA-MB-231, a metastatic TNBC cell line ^14^; and MCF-7, a luminal A cell line^15^) and of a non-tumorigenic breast cell, MCF-10A^16^. In addition, different signaling pathways related to antioxidant cell responses as well as regulated cell death induction have been investigated.

Collectively, our results showed that while MB-PDT is efficient in inducing multiple regulated necrosis mechanisms only in tumor cells, non-tumorigenic breast cell were able to mount an antioxidant response that led to impairment of the extensive photooxidative damage. We believe that these data highlight MB-PDT potential to be safe, accessible and an efficient adjunct to surgery for breast cancer treatment. Furthermore, this study contributes to the cancer and cell biology fields, providing further molecular mechanisms explaining why breast cells displaying distinct molecular makeups are able to undergo different regulated cell death pathways upon the same trigger.

## Results

### Human breast cells (BC) presenting variations in PDT sensitivity displayed differential cellular lipid composition

As a first step in our study, we confirmed previous results from our group^13^ by showing that cell death kinetics after MB-PDT exerted a higher impact in the malignant cell lines, being TNBC cells the most susceptible (**Figure 1A**). We then set out to investigate the molecular basis of these cell type specific differential responses to the therapy. Since MB-PDT relies on a massive intracellular generation of oxidant species^9,13,17^, with a widespread impact in membranes, we evaluated whether there was a link between the cellular lipid profile and the sensitivity to MB-PDT. For this purpose, we performed a comparative lipidomic profiling of BC using reversed-phase ultra-high-performance liquid chromatography (RP-UHPLC) coupled to electrospray ionization time-of-flight mass spectrometry (ESI-TOFMS). Around 487 different species were identified and classified in groups such as sphingolipids (SP), glycerophospholipids (GP), neutral lipids (NL), free fatty acids (FFA) and coenzyme Q (CoQ). The two first components of the Principal Component Analysis (PCA) explained 90.6% of the lipid content variance, showing a clear clustering between BC types (**Figure 1B**). Among all the obtained lipid classes, the main differences between BC were found on the amount of some species of NL [ex.: cholesteryl ester (CE), ceramide (Cer), diacylglycerol (DG) and triacylglycerol (TG)], coenzyme Q_10_ (CoQ_10_), and some species of GP [ex.: phosphatidylethanolamine (PE), phosphatidylinositol (PI), phosphatidylglycerol (PG), and phosphatidylcholine (PC), besides plasmanyl (o)- and plasmenyl (p)-GP, such as oPE, pPE, oPC and pPC (**Figure 1C and Supplementary Figure S1A, S1B**)]. All NL species were more abundant in breast cancer cells, mainly in TNBC, than in non-tumorigenic cells (**Supplementary Figure S1A**). Similarly to NL, a higher proportion of PI was found in malignant cells but in this case mainly in the luminal A subtype (**Figure 1C**). Moreover, while MDA-MB-231 and MCF-10A cells displayed similar levels of PE, the lowest levels of this class of lipids were observed in MCF-7 cells. Additionally, oPE and pPE were barely present in these cells (**Figure 1C**). Furthermore, considering all lipid species, MDA-MB-231 presented the highest levels of monounsaturated and polyunsaturated fatty acids (MUFAs and PUFAs, respectively) (**Figure 1D**), with special attention to arachidonic (ArA) and adrenic (AdrA) PUFAs (**Figure 1E and Supplementary Figure S1C**). Interestingly, MDA-MB-231 also presented the highest abundance of these acids esterified in PE (**Supplementary Figure S1D and S1E**). Worth of note that the non-tumorigenic cells displayed higher levels of CoQ_10_, compared to malignant cells (**Figure 1F**), suggesting that the better control of redox imbalance could play a role in the level of protection experienced by MCF-10 cells. Indeed, CoQ_10_ can trap lipophilic radicals and halt the propagation of phospholipid peroxidation^18^. These last observations led us to hypothesize whether ferroptosis, a form of regulated cell death (RCD), which is initiated by oxidative insults that occurs mainly towards PUFA, as ARA or AdrA^19^ (especially if they are esterified in PE^20^), would be a cell death mechanism induced by MB-PDT.

**Figure 1:**
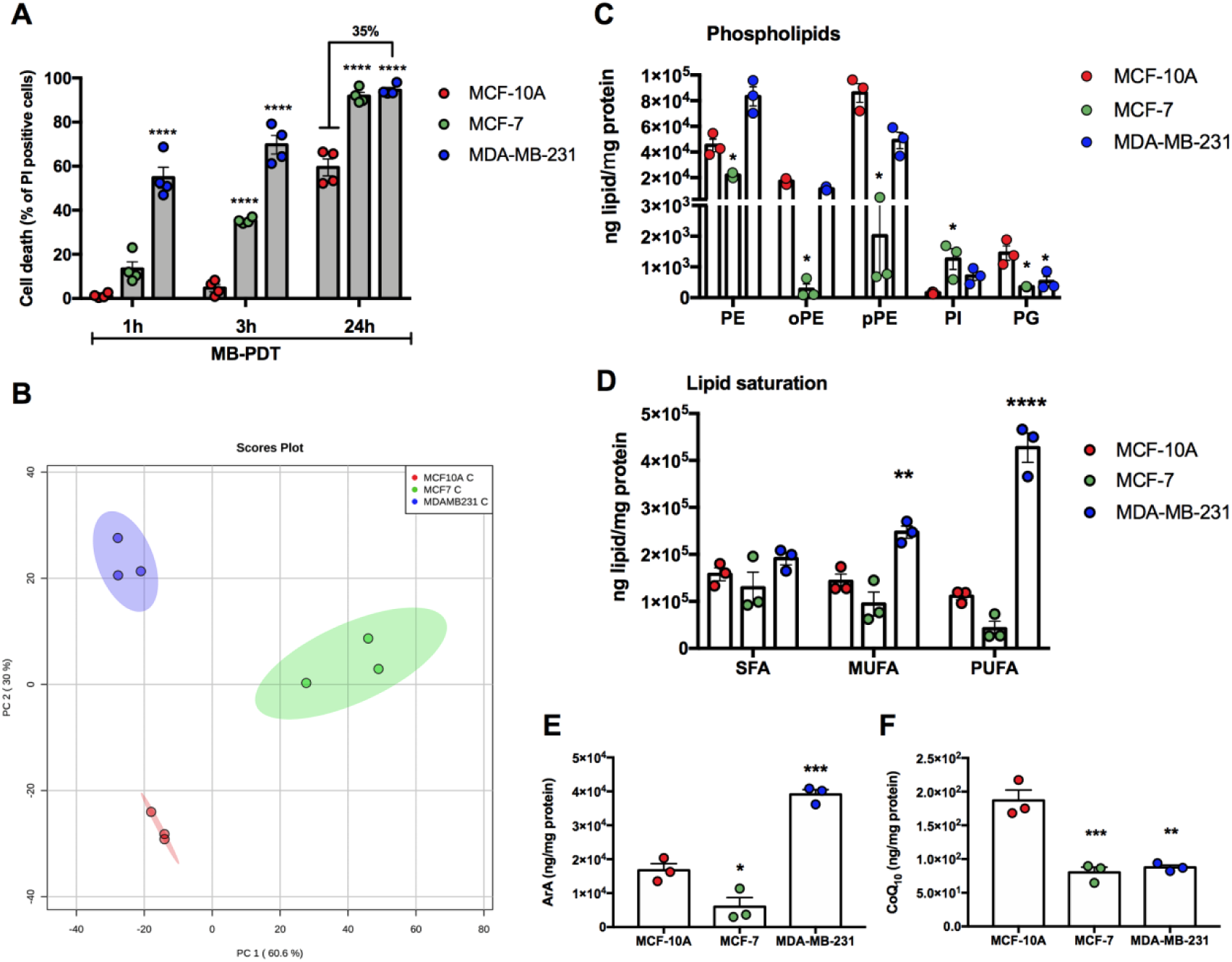
Human Breast Cells display differential sensitivity to MB-PDT and lipid composition. (A) Cell death induction after 1, 3 or 24 h of MB-PDT treatment. (B) Score plot of the Principal Component Analysis (PCA) of lipidomic data of BC. (C) Abundance of the glicerophospholipids: PE, oPE, pPE, PI and PG. (D) Abundance of lipids grouped by the degree of fatty acid chain saturation. Saturated fatty acid (SFA); monounsaturated fatty acid (MUFA); and polyunsaturated fatty acid (PUFA). (E) Abundance of ArA-containing lipids in BC. (F) Abundance of CoQ_10_. Dot color representation: MCF-10A in red; MCF-7 in green; MDA-MB-231 in blue. **** p<0.0001; *** p<0.001; ** p<0.005; *p<0.05 *vs* MCF-10A. Results are presented as mean ± S.E.M. Each dot represent an independent experiment. n≥3 independent experiments.

### MB-PDT induces ferroptosis in TNBC cells

As a prooxidant treatment, MB-PDT presents the potential to induce lipid oxidation and cell membranes damage^21^. We hence investigated whether MB-PDT would lead to the accumulation of oxidized lipids and could induce ferroptosis in BC. As a first step, the presence of the key components of ferroptosis ACSL4 (acyl-CoA synthetase long-chain family member 4) and GPX4 (gluthatione peroxidase 4) was evaluated in BC growing in basal conditions. Unlike MCF-7 cells, MCF-10A and MDA-MB-231 cells expressed ACSL4 (**Supplementary Figure 2**). Moreover, the non-tumorigenic cells presented higher protein levels of GPX4, compared to the luminal A or the TNBC cell lines (**Supplementary Figure 2**). The second step was studying whether cells submitted to MB-PDT undergo lipid peroxidation by using BODIPY-C11, a fluorescent probe for lipid oxidation. MCF-10A and MDA-MB-231 cells showed increased lipid peroxidation after MB-PDT treatment, while MCF-7 cells presented no differences (**Figure 2A, B**). Additionally, since iron available in the labile iron pools (LIP) has already been reported for being more prone to participate in ferroptosis, LIPs were analysed in the three cell lines before MB photooxidation. The highest LIP levels were found in MDA-MB-231 cells (**Figure 2C**). Remarkably, these data revealed that these cell lines constitute three distinct cell models to explore the role of ferroptosis in the context of photooxidation: one that exhibits PUFA but also disposes high lipid detoxification capacity, like GXP4 and CoQ_10_, (MCF-10A); another that besides displaying high levels of GPX4, does not posses high abundance of PUFA nor CoQ_10_ (MCF-7); and finally, the most agressive cell line containing a higher proportion of PUFA and LIP and a very low capacity to deal with lipid peroxidation due to the low levels of reduced gluthatione (GSH)^13^, GPX4 and CoQ_10_ (MDA-MB-231).

**Figure 2:**
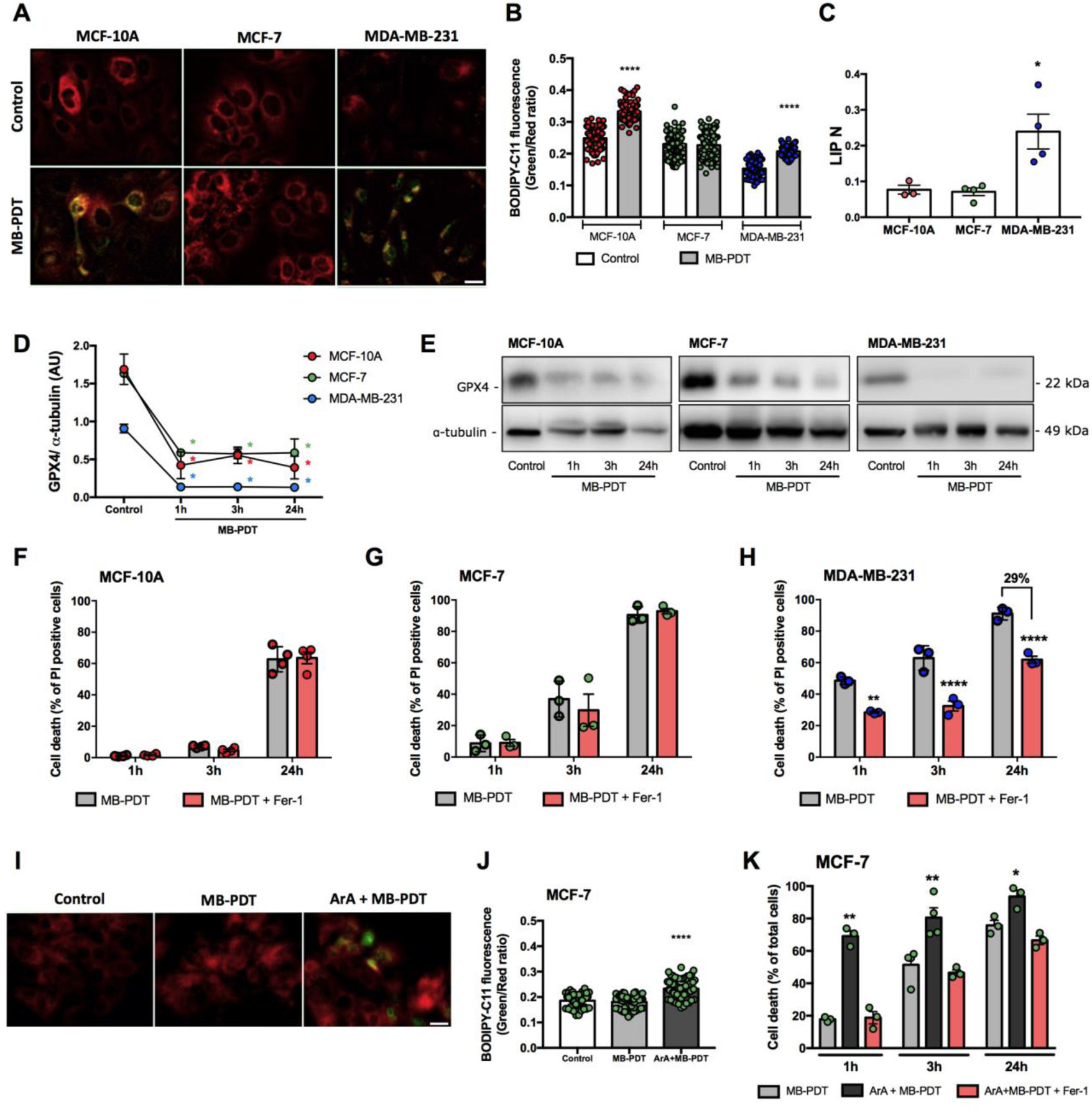
MB-PDT induces ferroptosis in PUFA- and LIP-enriched cells. (A) Representative images of lipid peroxidation in cells submitted or not to MB-PDT. Reduced (red) or oxidized (green) BODIPY-C11. (B) Graphical representation of the quantification of the oxidized/reduced BODIPY-C11 ratio/cell. Each dot represents an individual cell. Min. of 100 cells was analysed/experiment. **** p<0.0001 *vs* control of each cell line. (C) Intracellular labile iron poll (LIP), normalized by the total intracellular calcein incorporated per cell (LIP N). * p<0.05 *vs* MCF-10A. (D) GPX4 protein quantification in cells after 1, 3 and 24h of being submitted or not to MB-PDT (Control). * p<0.05 *vs* control of each cell line. (E) Representative images of Western blots of GPX4 of BC after being treated or not with MB-PDT, as indicated. Middle panels show the percentage of cell death after MB-PDT (1, 3 or 24h) in cells pretreated or not with Fer-1: (F) MCF-10A, (G) MCF-7 and (H) MDA-MB-231 **p<0.005 *vs* MB-PDT; ****p<0.0001 *vs* MB-PDT. (I) Representative images of lipid peroxidation in MCF-7 cells preincubated or not with ArA before being submitted or not to MB-PDT. (J) Corresponding quantification (as described in item B) of oxidized/reduced BODIPY-C11 ratio in MCF-7 cells. ****p <0.0001 *vs* control. (K) Cell-death percentage of MCF-7 cells pretreated or not with Fer-1 and/or ArA as indicated after 1, 3 or 24h of photooxidation induction. ** p<0.005 *vs* MB-PDT; *p<0.05 *vs* MB-PDT. n≥3 independent experiments. Dot color representation: MCF-10A in red; MCF-7 in green; MDA-MB-231 in blue. Results are presented as mean ± S.E.M.

In order to investigate the role of ferroptosis in these cells submitted to photooxidation, GPX4 protein levels were evaluated after MB-PDT treatment. Moreover, cells were pretreated with the ferroptosis inhibitor ferrostatin-1 (Fer-1), followed by MB-PDT and determination of cell death percentage. Results showed that despite the fact that MB-PDT promoted the depletion of GPX4 protein levels in all cell lines tested (**Figure 2D,E**), only MDA-MB-231 cells were protected by Fer-1 against photooxidation-induced cell death (**Figure 2F-H**). These results are consistent with our data indicating that MDA-MB-231 could have very low capacity to cope with lipid peroxidation and consequently would be more susceptible to ferroptosis.

To test whether the presence of PUFAs was required to induce ferroptosis and hence increase MB-PDT cytotoxicity, we pre-incubated MCF-7 with arachidonic acid (ArA) and then submitted the cells to MB-PDT. MCF-7 pretreated with ArA underwent lipid peroxidation after MB-PDT (**Figure 2I, J**) and became more sensitive to photooxidation (**Figure 2K**). Moreover, MB-PDT was now able to induce ferroptosis because Fer-1 significantly inhibited cell death (**Figure 2K**). These data demonstrated that lipid peroxidation was a cytotoxic insult triggered by MB-PDT and cells presenting low abundance of PUFAs were less affected. Furthermore, these results have indicated that in order to undergo ferroptosis in response to MB-PDT photooxidation, high levels of PUFA were required, as demonstrated for MDA-MB-231 and ArA-pretreated MCF-7 cells.

### Non-tumorigenic cells are more prone to mount an efficient antioxidant response against MB-PDT

In order to further understand the resistance of MB-PDT-induced cell death observed in non-tumorigenic cells, the basal levels of other key antioxidant-related proteins, such as NF-E2-related factor 2 (NRF2), glucose 6-phosphate dehidrogenase (G6PD), copper/zinc and manganese superoxide dismutases (SOD1 and SOD2), and glutathione reductase and synthetase (GR and GS) were analysed and compared. Luminal A tumor-derived cells displayed the highest basal protein levels of G6PD, SOD1, SOD2 (**Supplementary Figure 3A-F**). Interestingly, the expression of GS in both tumor cells was extremely low compared to MCF-10A cells (**Supplementary Figure 3A, G**). These data indicated that tumorigenic cells possess less capacity to synthesise new GSH molecules and that, among all the models tested, TNBC cells would be potentially more susceptible to pro-oxidative damage^13^.

**Figure 3:**
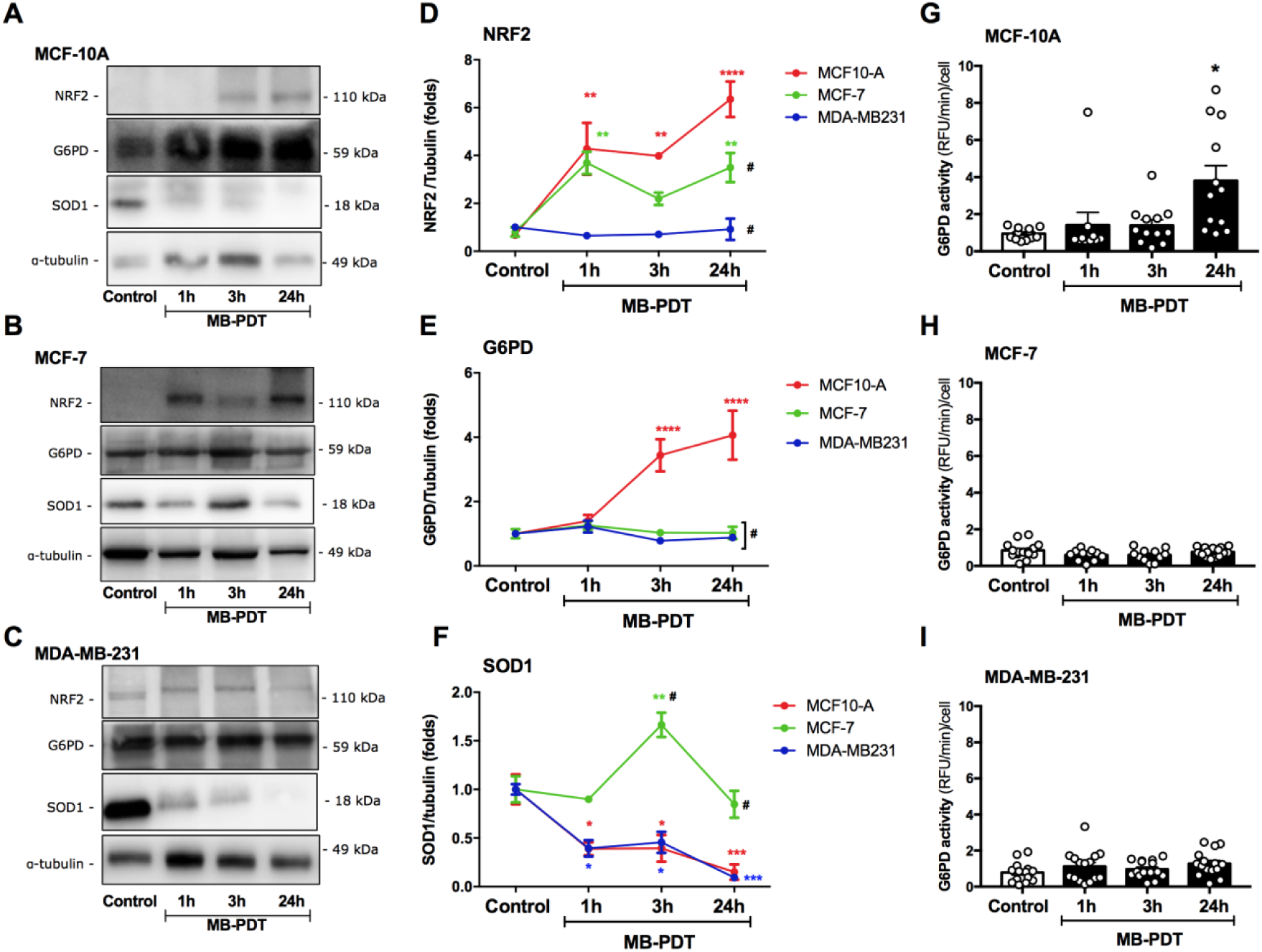
Non-tumorigenic breast cells display antioxidant response to MB. Representative images of Western blots of NRF2, G6PD and SOD1 for each BC after being submitted or not with MB-PDT: (A) MCF-10A, (B) MCF-7 and (C)MDA-MB-231. Western blot quantifications of (D) NRF2, (E) G6PD and (F) SOD1 of each BC treated or not with MB-PDT (PDT). Results are presented as folds *vs* Control condition. Color representation: MCF-10A in red; MCF-7 in green; MDA-MB-231 in blue. G6PD activity in (G) MCF-10A, (H) MCF-7 and (I) MDA-MB-231 cells after being submitted or not to MB-PDT (PDT). n≥3 independent experiments. **** p<0.0001; *** p<0.001; ** p<0.005; *p<0.05 *vs* Control; # p<0.05 *vs* MCF-10A at each time point. Results are presented as mean ± S.E.M.

To further explore the antioxidant response of these cell lines to MB-PDT, we analysed the protein levels of some of these sets of enzymes in cells submitted or not to photooxidation. Despite the lack of differences in basal NRF2’s levels between cells (**Supplementary Figure 3A,E**), upon MB-PDT this transcription factor was significantly increased in MCF-10A, slightly increased in MCF-7 and not modulated in MDA-MB-231 cells (**Figure 3A-C,D**). Intriguingly, NRF2 rise was accompanied by higher G6PD protein levels (**Figure 3A-C,E**) and activity (**Figure 3G-I**) only in the non-tumorigenic cells. While cellular levels of SOD1 significantly peaked at 3h after MB-PDT and then remained high up to the last time point in MCF-7 cells, a pronounced depletion of SOD1 protein was observed early in both MCF-10A and MDA-MB-231 cells (**Figure 3A-C,F**) submitted to the treatment. These observations may indicate that cytosolic SOD could play a protective role in MCF-7 cells submitted to photooxidation, highlighting that a differential cellular response mechanism against MB-PDT has been triggered.

Altogether, these results allowed us to conclude that the non-tumorigenic cells were able to activate a proficient antioxidant response through the increase of G6PD, which in turn would lead to the higher production of NADPH that could then be used to regenerate GSH, as well as reduced CoQ_10_, contributing to maximize the detoxification process and thus minimize cell death.

### MB-PDT can also trigger necroptosis

In order to investigate whether necroptosis was a cell death mechanism activated by MB-PDT in BC, the basal protein levels of key components of this pathway, such as RIPK1, RIPK3 and MLKL (Receptor Interacting Protein Kinases-1 and -3, and Mixed Lineage Kinase domain-Like protein, respectively), were first checked. MCF-7 cells displayed he highest levels of all the proteins analysed (**Supplementary Figure 4A-D**). Activation of this pathway was next assessed by monitoring MLKL phosphorylation levels (pMLKL). Interestingly, while pMLKL levels increased in all cancer cells submitted to MB-PDT, no sign of necroptosis activation was observed in the non-tumorigenic cells (**Figure 4A-C**). The role of necroptosis in MD-PDT cell death was further investigated in BC pretreated with necrostatin-1 (Nec-1) or necrosulfonamide (NSA), RIPK1 and MLKL inhibitors respectively. Our data showed that only the tumorigenic cells were protected from MB-PDT effects in the presence of the necroptosis inhibitors Nec-1 or NSA (**Figure 2D-F**). Moreover, the relevance of necroptosis as a cell death mechanism induced by MB-PDT in breast tumor cells was also confirmed by silencing the expression of RIPK3 or MLKL (**Supplementary Figure 4E-H**). Altogether, these data indicated that MB-PDT is capable of activating necroptosis in both luminal A and TNBC cells. However, up to this point of the study, the cytotoxic mechanism triggered by MB-PDT in the non-tumorigenic cells was still not elucidated.

**Figure 4:**
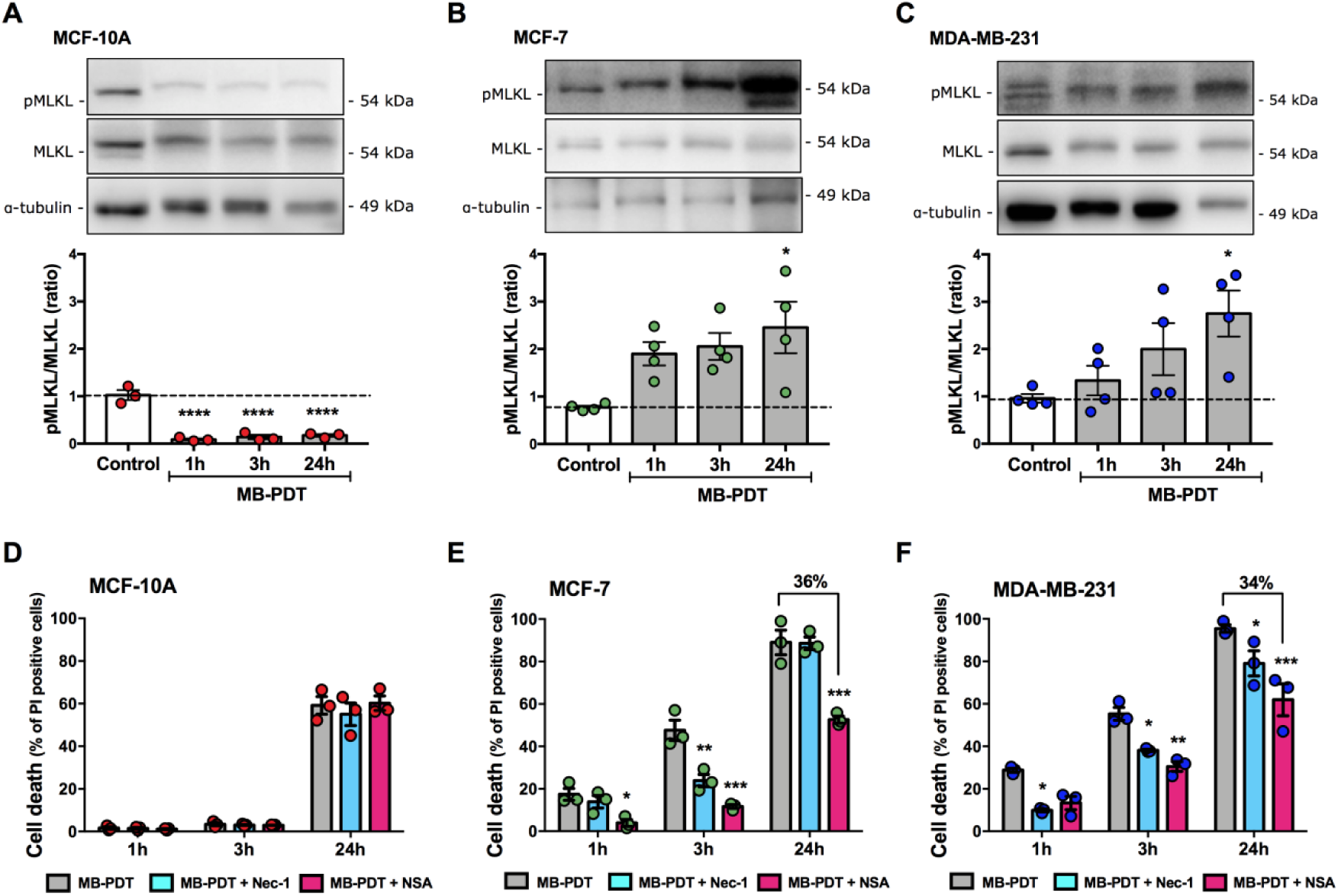
MB-PDT activates necroptosis in tumorigenic breast cells. (A) MCF-10A; (B) MCF-7 and (C) MDA-MB-231 cells were submitted or not to MB-PDT. 1, 3 and 24h after irradiation, cells were lysed, and protein extracts were submitted to Western blot. Representative images of Western blots of pMLKL, MLKL and tubulin and the corresponding quantification of pMLKL/MLKL ratio are shown. *p<0.05 *vs* Control (n=4 independent experiments). (D) MCF-10A, (E) MCF-7 and (F) MDA-MB-231 were pretreated or not with Nec-1 or NSA and then submitted or not to MB-PDT. Cell death was evaluated 1, 3 or 24h after irradiation. *** p<0.001; ** p<0.005; *p<0.05 *vs* MB-PDT. Dot colors representation: MCF-10A in red; MCF-7 in green; MDA-MB-231 in blue. Results are presented as mean ± S.E.M (n=3 independent experiments).

### MB-PDT induces lysosome damage

Based on the fact that we had previously reported that MB was mainly accumulated in lysosomes^13^, the hypothesis that photoactive MB could be capable to damage lysosome membrane and thus induce lysosome-dependent cell death (LDCD) was raised and tested. The first evidence was obtained by assessing the cytosolic activity of the lysosomal cathepsin B. In fact, MB-PDT induced lysosomal membrane permeabilization (LMP) (**Figure 5A**). The involvement of LDCD mediating the PDT effects by inhbiting cathepsin B with a small molecule inhibitor (CA-074) was then investigated. Cell incubation with this inhibitor was able to significantly decrease MB-PDT cytotoxicity (**Figure 5B-D**). Moreover, the results have also indicated that LMP is a common event triggered by MB-PDT in all BC analysed. Additionaly, since the levels of pMLKL were decreased upon cathepsin B inhibition, the results pointed at a yet undescribed possible cross-talk between LDCD and necroptosis induction after photooxidation (**Figure 5E,F**).

**Figure 5:**
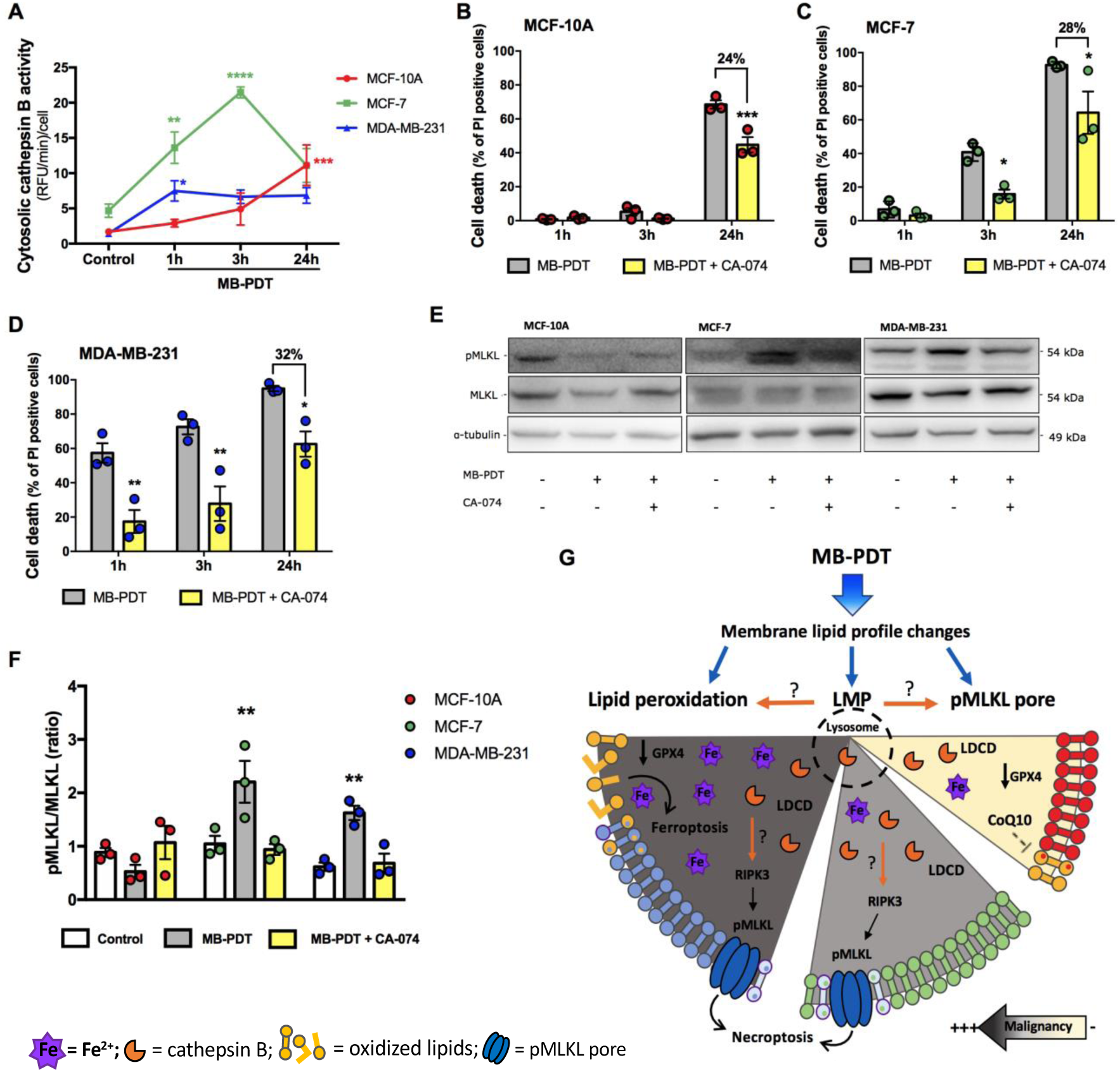
MB-PDT-induced LMP is involved on cell death of breast cells. (A) Cells were submitted or not to MB-PDT and cytosolic cathepsin B activity was analysed after 1, 3 or 24h of cell treatment. **** p<0.0001; *** p<0.001; ** p<0.005; *p<0.05 *vs* Control. (B) MCF-10A, (C) MCF-7 and (D) MDA-MB-231 cells were pretreated or not with CA-074 and then submitted or not to MB-PDT. Cell death was analysed after 1, 3 or 24h after cell irradiation: *** p<0.001; ** p<0.005; *p<0.05 *vs* MB-PDT. (E) Representative images of Western blots of pMLKL, MLKL and tubulin of BC treated or not with CA-074 and MB-PDT after 24h, as indicated. (F) pMLKL/MLKL ratio of cells after 24h they were treated or not with CA-074 and then submitted to MB-PDT. ** p<0.005; *vs* Control. Dot colors representation: MCF-10A in red; MCF-7 in green; MDA-MB-231 in blue. n≥3 independent experiments. Graph results are presented as mean ± S.E.M. (G) Cell death mechanisms activated by MB-PDT in BC ranging from non-malignant to very aggressive tumorigenic cells (from right to left). Upper part shows that photooxidation induces membrane lipid profile changes such as lipid peroxidation, LMP and/or pMLKL pore formation. Bottom part of the figure represents the differential lipid composition of each BC analysed (represented by different lipid membrane colors) and which mechanisms are activated in each one (LDCD: lysosome dependent cell death, necroptosis or ferroptosis). MDA-MB-231 cells: dark gray background, with higher proportion of lipids to undergo lipid-peroxidation; MCF-7: gray background displaying lipids not susceptible to peroxidation; MCF-10A: yellow background and bearing intermediate abundance of lipids being susceptible to oxidation.

We demonstrated that MB-PDT trigger multiple RCD in BC by inducing modifications in lipid membranes. In malignant cells, our data pointed that LMP was followed by MLKL phosphorylation and necroptosis. In case of high PUFA-lipid content, in addition to LDCD and necroptosis, cells were also able to undergo ferroptosis, which was triggered via lipid peroxidation, GPX4 depletion and failure of the cell capacity to activate the cell responses involved in oxidative damage detoxification. Moreover, in these cells, oxidative chain reactions could be facilitated and amplified by higher concentration of labile iron intracellular pool, resulting in a massive and faster cell death (**Figure 5G**).

## Discussion

Non-tumorigenic human breast epithelial cells were more resistant to PDT than their malignant and more metastatic counterparts^13^. Up to date, there are no studies addressing a possible association of a defined lipid profile with aggressiveness and susceptibility to undergo different cell death subroutines upon massive oxidant species generation^22,23^. Therefore, the present study has focused on understanding the molecular pathways underlining these different cell behaviors upon photooxidative damage (**Figure 6**). We have demonstrated that MB-PDT can activate multiple cell death mechanisms, namely ferroptosis, necroptosis and LDCD. This study has also assessed the cellular makeups in lipid composition and antioxidant machinery, which underlie the unique capacity of the most aggressive breast cancer cells tested to undergo ferroptosis upon MB-PDT. Moreover, our data have further reinforced the potential of this therapy to present fewer side effects on non-cancerous breast tissue, by providing several evidences on how PhOxS triggered by MB-PDT has barely affected antioxidant capacity of non-tumorigenic breast cells to deal with oxidative damage (**Figure 6**).

**Figure 6:**
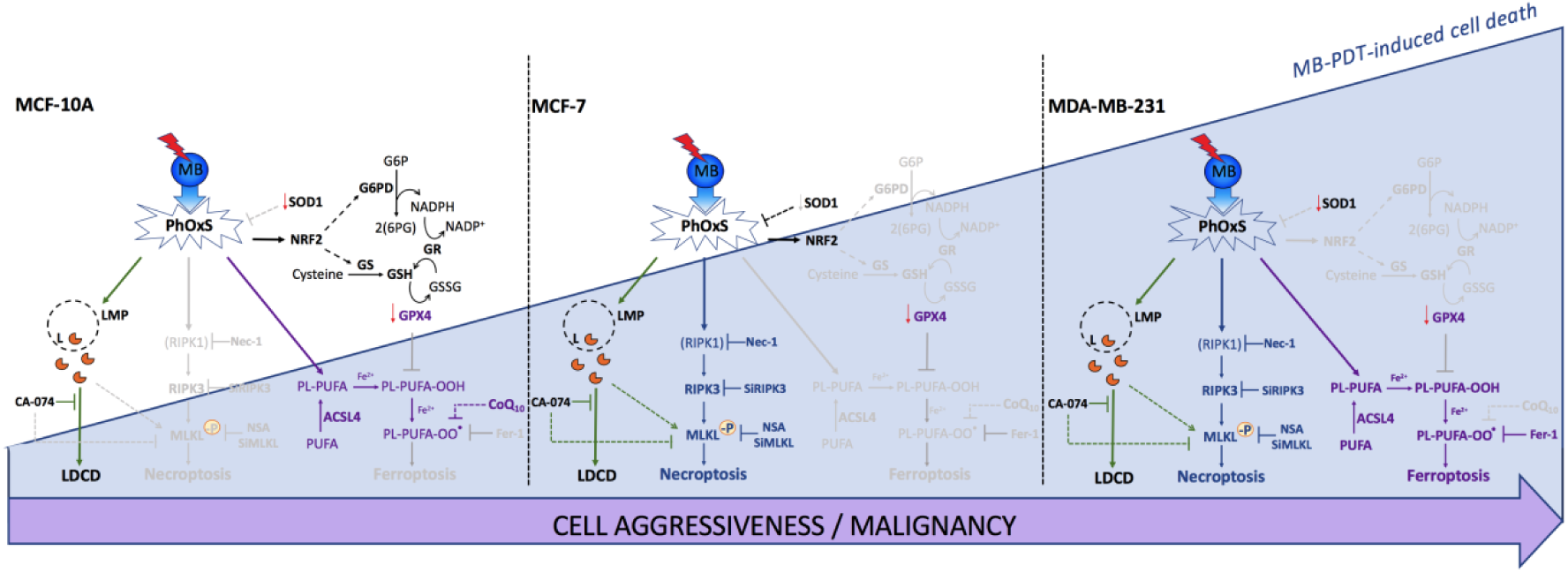
Scheme summarizing the antioxidant and cell death mechanisms activated in human breast cells by MB-PDT. Breast cells display different susceptibility to photooxidative stress (PhOxS) induced by MB-PDT, being the highest effect observed in MDA-MB-231. In this cell line, no antioxidant response was mounted upon PhOxS. In addition, low levels of GPX4 and CoQ_10_, combined with high amount of iron (Fe^2+^) and PUFA-phospholipid content (PL-PUFA), resulted in ferroptosis activation by MB-PDT (purple arrows and letters). This cell death was inhibited by Fer-1 pre-treatment. Lysosomal damage was observed in all cell lines, evidenced by the release of cathepsin B through lysosomal membrane permeabilization, LMP, (green arrows and letters). Pre-treatment with CA-074, a cathepsin B inhibitor, alleviated cell death. In both tumorigenic cells, MDA-MB-231 and MCF-7, necroptosis activation with MLKL plasma membrane pore formation (blue arrows and letters) was observed. Inhibition of RIPK1, RIPK3 or MLKL phosphorylation, by gene silencing or pre-treatment with Nec-1 or NSA, rescue tumorigenic cells from death. A possible link between LMP and necroptosis was found in tumorigenic cells (green dotted arrows). Because MCF-7 cells lack significant amounts of oxidizable phospholipids, lipid peroxidation was not observed and therefore ferroptosis did not contribute to death. However, a complete antioxidant response was not sustained in these cells, making them also highly affected by MB-PDT. The scenario after PhOxS was quite different for MCF-10A cells. Even occurring LMP and lipid peroxidation, they were significantly more resistant to MB-PDT than the other cells. Neither ferroptosis nor MLKL phosphorylation and necroptosis were observed.

By using high resolution lipidomics, we mapped the lipid content in three different BC models displaying different susceptibilities to PDT-induced photooxidative redox imbalance. In the recent years lipids have emerged as key regulators of cell fate^24,25^. Increasing amount of evidences have highlighted their functions as triggers, executors or modulators of plasma membrane components that act as platforms for RCD execution^26^. Lipids, displaying different susceptibilities to undergo chemical modifications can execute their functions on cell death subroutines by modulating membrane physicochemical properties^27,28^. For example, exposure of lipids to sources of free radicals, molecular oxygen and redox-active metal ions, such as low-valent iron, results in lipid peroxidation^29^. Noteworthy, in the context of PDT the direct contact within lipid membranes and the excited triplet state of the PS molecule, during photosensitized oxidations, also results in membrane peroxidation^8^. We provided strong evidences of membrane lipid peroxidation in cells containing high proportion of PUFAs submitted to MB-PDT-induced photooxidation. The presence of methylene groups flanked by carbon-carbon double bonds make PUFAs particularly good substrates for oxidations^30^. Therefore, the subsequent accumulation of lipid peroxidation products culminates in the permeabilization of the membrane^31^.

Other groups have already correlated an increased abundance of PUFAs with ferroptosis induction^20,32^. Moreover, the mechanism of ferroptosis was identified as specifically dependent on intracellular iron availability and not on other metals^33^. Herein, on one hand, it has been shown that highly aggressive breast cancer cells presented the highest PUFAs and LIP contents and consequently were the only cells undergoing ferroptosis upon MB-PDT. On the other hand, we were able to demonstrate that exogenous supplementation of ArA was sufficient to bypass the protective effect conferred by the lack of ACSL4 in the non-invasive MCF-7 cells submitted to MB-PDT. ACSL4 is the acyl transferase that catalyzes the conversion of long-chain fatty acids, preferentially arachidonate and eicosapentaenoate, to their active form acyl-CoA in PUFA-containing lipid synthesis^34,35^. These results pointed PUFAs as key lipid modulators involved in the molecular mechanisms of MB-PDT cytotoxicity. Exogenous metabolites, including lipids, have been reported to be potent modulators of cell function and fate and could conceivably be relevant in several contexts *in vivo*, especially for cells that can extract exogenous lipids directly from the bloodstream^36,37^. In healthy individuals, raising serum PUFA levels could provide a means to improve the lethality of existing pro-ferroptotic agents against cancer cells^37^. Moreover, our studies have suggested that the administration of exogenous PUFAs before the photooxidation could potentially help to improve the power of this therapy by tackling tumor cell death resistance to MB-PDT.

In physiological conditions, lipid peroxides are reduced to non-toxic lipid alcohols by GPX4, at the cost of the oxidation of two molecules of GSH per lipid molecule^38^. However, this enzyme is absent or inactive during ferroptosis, resulting into toxic lipid accumulation^39,40^. As a consequence, the inhibition of GPX4 or depletion of GSH has emerged as therapeutic strategies to induce cancer cell death^41^. We have previously shown that MB-PDT was able to deplete GSH levels in breast tumor cells^9^. Additionally, in the present study the levels of GPX4 were decreased in all BC tested upon photooxidation, underscoring MB-PDT as a potential ferroptosis inducer. This observation is consistent with the fact that sensitivity to GPX4 inhibitors varies greatly across cell lines and ferroptosis may have additional regulation mechanisms^42^. Indeed, a glutathione-independent regulation axis involving the ferroptosis suppressor protein 1 (FSP1), previously called apoptosis-inducing factor mitochondrial 2 (AIFM2), was recently uncovered^18,43^. FSP1 acts as a NAD(P)H-oxidoreductase that reduces coenzyme Q-10 (CoQ_10_), which can, in turn, trap lipophilic radicals and halt the propagation of lipid peroxidation^18,43^. Interestingly, our results have shown that non-tumor breast cells presented the highest abundance of CoQ_10_ among the three cell lines and were able to survive despite undergoing lipid peroxidation and GPX4 depletion. CoQ_10_ can be reduced in the presence of NADPH and breast non-tumorigenic cells were the only model where it was observed an increase of activity of a key enzyme for the generation of NADPH cellular supply, G6PD, upon MB-PDT. Therefore, one can speculate that the FSP1–CoQ_10_–NAD(P)H axis could be operating to suppress phospholipid peroxidation and ferroptosis after photooxidative damage in the non-tumorigenic breast cells. It is important to note that this particular effect could be associated with the low side effects of this therapy.

Previous results from our group have demonstrated the importance of optimizing fluence rates in order to provide exhaustion of the cell antioxidant responses to circumvent therapy resistance of breast tumors using MB-PDT^9^. We now observed that the basal levels of key antioxidant proteins were lower in TNBC cells, further extending the knowledge that these cells exhibit significant metabolic alterations compared to luminal tumors^13,44^. Moreover, our data indicated that despite the non-invasive MCF-7 cells displayed high basal levels of most of the antioxidant proteins evaluated, they were not able to mount such an efficient antioxidant response against photooxidation when compared to the one displayed by the non-tumorigenic breast cell line, which were more resistant to PDT.

As a prooxidant multifaceted therapy, it is difficult to precisely address where the trigger of PDT will be. Since in general the first photoreaction will occur in the vicinity of where the PS presents the highest concentration^45^, determining its subcellular localization could give us a clue. We have previously shown that in these cell lines MB accumulates preferentially in the lysosomes^13^. Several reports have suggested that lysosome integrity is critical to maintain cellular homeostasis, because it ensures the proper localization of lysosomal enzymes and organelle function. Upon damage, lysosomal content leaks into the cytoplasm as a consequence of lysosome membrane permeabilization (LMP)^46^. In the present study we identified the occurrence of MB-PDT-dependent LMP. In principle, this process should represent little danger, because most lysosomal proteases are inactive at neutral pH. However, some cathepsins, such as cathepsins B, D, and L, remain active at neutral pH and can trigger a cascade of events, including the proteolytic modification of molecules implicated in cell death pathways^47^. Therefore, LMP can be followed by a multi-pathway process that results in LDCD, which, in most cases, can be prevented by inhibiting lysosomal protease activity^46,48^. Interestingly, LDCD was the only PDT-induced process contributing to the low cell death observed in non-tumorigenic breast cells submitted to MB-PDT.

Cytosolic cathepsins usually lead to apoptotic RCD by catalyzing the proteolytic activation or inactivation of several substrates, including BAX and anti-apoptotic BCL2 family members^48^. However, we have previously shown that apoptosis was not the main cell death mechanism activated in BC submitted to MB-PDT^13^. In line with this, it has been recently demonstrated, indeed, that LDCD does not necessarily manifest itself with an apoptotic morphotype and an intriguing connection is emerging between LMP, the adaptive responses to stress, and other RCD subroutines^49,50^. For example, lysosomes have an essential role in autophagy and cellular iron homeostasis, being a major source of free iron due to the degradation of ferritin in a process called ferritinophagy^51–53^. Thus, lysosomal damage could allow this metal to be more bioavailable to peroxidation reactions. Therefore, upon lipid peroxidation and in the absence of a proper detoxifying response, as occurred in breast tumorigenic cells upon MB-PDT, LMP may facilitate catalysis of iron-dependent reactions and increase ferroptosis susceptibility.

Based on our data that ferroptosis only occurs in TNBC cells and that its inhibition did not completely rescue cells from death, we hypothesized that MB-PDT was activating more than one regulated necrotic pathway simultaneously. Intriguingly, we have also detected activation of necroptosis only in tumorigenic cells submitted to photooxidation damage. Necroptosis is triggered by perturbations of extracellular or intracellular homeostasis that critically depend on the kinase activities of RIPK1, at least in some settings, RIPK3, and consequent phosphorylation, oligomerization and migration of MLKL (mixed lineage kinase domain-like protein) to the plasma membrane^3,48^. MLKL oligomers form channels on the plasma membrane, leading to high osmotic pressure, water influx, release of intracellular components, and eventual plasma membrane rupture^54^. To undergo necroptosis, MLKL engagement requires the presence of specific lipids at the plasmatic membrane. Phosphatidyl inositol phosphates including, phosphatidylinositol-5-phosphate and phosphatidylinositol-4,5-bisphosphate have been described to function as lipid receptors of MLKL in the inner leaflet of the plasma membrane^55–57^. These lipids are products of kinases that can phosphorylate the 3-, 4-, and 5-hydroxyl groups of the inositol head group of PI lipids^58^. Our study revealed that both breast cancer cell lines presented higher levels of overall PI, compared to the non-tumorigenic cells. Consistent with this observation, only breast cancer cells underwent MLKL phosphorylation and necroptosis. It is thus reasonable to suppose that lipid composition of normal cells does not sustain the necroptosis membrane pore formation, also contributing to their resistance to photooxidation-induced cell death.

It has been previously shown by others that RIPK1 and RIPK3 can be degraded in lysosomes and that inhibition of lysosomal function, with LMP, leads to this kinases accumulation and necroptosis induction^59,60^, strengthening the possibility of a link between LMP and necroptosis. Indeed, we have shown that, in breast cancer cells submitted to MB-PDT, lysosomal cathepsin inhibition was able to suppress cell death and MLKL phosphorylation, providing a clear evidence of the existence of cross-talks between LDCD and necroptosis, triggered by MB-PDT. We suggest that, because as MB localizes mainly in lysosomes of these cells, LMP is an inevitable consequence of MB photoactivation, and thus may be driving the RCD events. The activation of different cell death subroutines will then depend on the availability of the required components of each pathway. We have described that different human breast epithelial cells, from non-tumorigenic to very aggressive malignant cells, display distinct structural and metabolic traits within different signaling pathways were activated upon MB-PDT. In this study, LMP appears as the common event, which was then followed by an antioxidant response (in non-tumorigenic cells), necroptosis (in non-invasive tumor cells) or both necroptosis and ferroptosis (in highly aggressive tumor cells).

Collectively, our data have provided molecular mechanisms behind a hitherto unexplored therapeutic approach, which have simultaneously activated alternative tumor regulated cell death pathways while preserving the integrity of most of the non-tumorigenic cells. This fact is of fundamental importance since despite all the recent technological improvements, breast cancer still has significantly impacts on global health, being disease recurrence and metastasis the bottleneck for an effective clinical treatment^2,3^. This study contributes to a better understanding of breast cancer susceptibility to photooxidation-induced damage. Our results could provide the rational and know-how needed to maximize the therapeutic clinical application gain of MB-PDT.

## Materials and methods

### Cell culture

MCF-10A (ATCC CRL-10317(tm)) cells were maintained in phenol red-free DMEM-F12 (Sigma-Aldrich, St. Louis, MO, USA) supplemented with 5% heat-inactivated horse serum (Thermo Scientific, Waltham, MA, USA), insulin (10 μg/mL; Sigma-Aldrich), cortisol (500 ng/mL; Sigma-Aldrich), cholera enterotoxin (100 ng/mL; Sigma-Aldrich), and epidermal growth factor (20 ng/mL; Sigma-Aldrich). MCF-7 (ATCC HTB-22(tm)) cells were maintained in phenol red-free DMEM-F12 (Sigma-Aldrich) with 10% heat-inactivated FBS (Vitrocell Embriolife, Campinas, Sao Paulo, Brazil). MDA-MB-231 (ATCC HTB-26(tm)) cells were cultured in phenol red-free RPMI 1640 (Sigma-Aldrich) with 10% heat-inactivated FBS (Vitrocell Embriolife). All cultures were maintained at 37°C under water-saturated atmosphere containing 5% CO_2_.

### Photodynamic treatment (MB-PDT)

An aqueous solution containing of Methylene blue (MB, Labsynth Products, São Paulo, Brazil) was used as PS. Cells were seeded at 30.000 cells.cm^-2^ and after 24 h they were incubated for 2 h with 20 μM of MB solution, in phenol red-free medium and maintained in these conditions during both irradiation and post-treatment time. All assays were performed with a reduction of 75% of medium supplements. The whole microplate was irradiated from the top with a light-emitting diode (LED) array, with maximum emission wavelength at 640 nm. The irradiation time was 16 min with a total light dose was 4.5 J.cm^-2^ (fluence rate of 4.7 mW.cm^-2^). Control experiments, such as cells exposure or not to the PS or exposure or not to light were performed. For cell death inhibition assays, inhibitors were pre-incubated with MB solution 2 h before irradiation: Fer-1 (Cayman Chemical, Ann Arbor, Michigan, EUA, 1 µM); Nec-1 (Sigma-Aldrich, 10 µM); NSA (abcam, Cambridge, United Kingdom, 5 µM); or CA-074 (Millipore, Burlington, Massachusetts, EUA, 400 nM). Pretreatment with ArA (Sigma-Aldrich, 24 µM) was performed 16 h before MB incubation.

### Cell death evaluation

After treatments cells were stained with the DNA-binding dyes Propidium iodide (PI, Sigma-Aldrich) and Hoechst 33342 (HO, Sigma-Aldrich) for 10 min. Following incubation, the percentage of viable and dead cells was determined using an inverted fluorescence microscope (Nikon Eclipse Ti, Kyoto, Japan) with 20x of magnification. Fluorescence of labelled cells was detected using laser 461 nm and 545 nm for excitation of HO and PI respectively. The cultures were evaluated according to: the total number of cells, determined by counting the nuclei stained with HO; and the number of dead cells determined by the number of nuclei stained with PI. A minimum of 500 cells was counted in each experimental condition. Results were expressed as percentage of dead cells.

### Transient oligonucleotide transfection

The transfection was performed 24 h after seeding of 1.5×10^4^ cells.cm^-2^. The lipid carrier Lipofectamine RNAiMAX (Life Technologies, Carlsbad, CA, USA) and the Silencer® Select pre-designed siRNA (Life Technologies) for human RIPK3 (GGCAAGUCUGGAUAACGAAtt) or for human MLKL (CCCGUUUCAAGGUGAAGAAtt) were used. “AllStars negative control siRNA” (Qiagen, Venlo, Netherlands) was used as a negative siRNA control of scrambled sequence (siControl). Lipid-RNA complexes were formed in Opti-MEM (Invitrogen) in a proportion of 0.6 μL of Lipofectamine to 0.45 μL of 20 μM siRNA, at room temperature for 20 min and were further added to cells in antibiotic-free medium to reach a final volume of 300 μL (in a 24-well plate) for overnight transfection. Cells were maintained in culture for a 24 h recovery period before experiments were carried out. The efficiency of transfection/silencing was validated by Western blot.

### Western blots

Cells were lysed in RIPA Buffer (Sigma-Aldrich) containing protease inhibitor (Roche, Basel, Switzerland) and phosphatase inhibitor (Sigma-Aldrich) cocktails. 30 μg protein were loaded on a 12% denaturing gel and proteins were separated by SDS-PAGE. Proteins were transferred by tank blot onto a PVDF membrane that was subsequently blocked in a solution containing 5% Blocking Buffer (Life Technologies) and 5% BSA (1:1) at 4°C overnight. Primary antibodies were diluted in a solution of 5% BSA in PBS (**Table 1**) and were incubated overnight at 4°C. Membranes were washed three times in PBS-Tween (0.1%) and then incubated at RT for 1h with HRP-labeled secondary antibodies, diluted in a solution of 1% BSA in PBS. The protein-antibody complex was visualized by using enhanced chemiluminescence (Millipore Corporation, Billerica, MA, USA). Images were acquired using Uvitec Image System (Cleaver Scientific Limited, Cambridge, UK). Quantitative densitometry was carried out using the ImageJ software (National Institutes of Health). The volume density of the chemiluminescent bands was calculated as integrated optical density × mm^2^ using ImageJ Fiji.

**Table 1:** Antibodies

| *Protein* | *Company* | *Catalog* | *Dilution* |
| --- | --- | --- | --- |
| RIPK1 | BD | #610458 | 1:1000 |
| RIPK3 | Cell Signaling | #13526 | 1:1000 |
| phospho(S345) MLKL | abcam | ab196436 | 1:1000 |
| MLKL | abcam | ab184718 | 1:1000 |
| ACSL4 | Santa Cruz | sc-271800 | 1:200 |
| GPX4 | abcam | ab125066 | 1:1000 |
| G6PD | abcam | ab993 | 1:2000 |
| NRF2 | abcam | ab137550 | 1:1000 |
| SOD1 | abcam | ab51254 | 1:2000 |
| SOD2 | abcam | ab16956 | 1:1000 |
| Glutathione Reductase | abcam | ab128933 | 1:2000 |
| Glutathione Synthetase | abcam | ab133592 | 1:2000 |
| alpha-tubulin clone B-5-1-2 | Sigma-Aldrich | T5168 | 1:10.000 |
| anti-rabbit | Vector Laboratories | PI1000 | 1:1000 |
| anti-mouse | Vector Laboratories | PI2000 | 1:1000 |

### Cathepsin activity assay

Cells were washed with phosphate buffer (PBSA: NaCl 137 mM, KCl 2.7 mM, Na_2_HPO_4_ 10 mM, KH_2_PO_4_ 1.8 mM, pH 7.2) and detached from the plate using trypsin solution (0.5% p/v). The cells were then centrifuged at 800 x g for 2 minutes. Cell pellets were washed with PBSA and resuspended in 2 mL PBSA. The number of cells was counting in a hemocytometer. Samples were then homogenized in syringes with insulin needle 10 times, and centrifuged at 4°C, 700 x g for 10 minutes. The supernatants were collected and centrifuged at 4°C, 25000 x g for 2 hours for cytosol and organelles fractionation. The supernatants (cytosolic fraction) were used in cathepsin-B/L kinetics assays using Z-FR-MCA as substrate (10 µM) in 100 mM citrate phosphate buffer, pH 6. Protease activity was evaluated at an excitation wavelength of 380 nm and an emission wavelength of 460 nm using a 96-well plate in a spectrofluorometer (SpectraMAX M2, Molecular Devices, Sunnyvale, CA, USA). Fluorescence intensity values were collected every 5-minute intervals for 1 hour. Activity units were calculated as: [relative fluorescence units (RFU)/min]/number of cells. Each experiment was performed in duplicate. At least three independent experiments were performed for each cell type and condition.

### G6PD activity assay

The determination of G6PD activity was performed as already described^61^. Reaction medium was incubated at 37°C. Fluorescence intensity values were collected every 2 min during 1h at 500 nm in a SpectraMax M2 microplate reader (Molecular Devices). Activity units were calculated as: (RFU/min)/number of cells. Each experiment was performed in duplicates and at least three independent experiments were performed for each cell type and condition.

### Lipid peroxidation analysis

After irradiation, the cells were incubated with 1 µM BODIPY C11 at 37°C during 20 min. The probe was then removed and 1 mL of fresh medium without serum and phenol red was added. Cells were imaged using a fully motorized Leica DMi8 widefield microscope (from Leica Microsystems) using the FITC and Texas Red filter sets and a 20x objective. Imaging was performed on two independent biological replicates. In each independent experiment at least 4 different images (100 cells) per condition were analysed. All imaging acquisition parameters were kept constant for each experiment. Images were quantified using ImageJ Fiji and quantified as follows. Cell outlines were free-handed drawn on the bright field channel to generate a cell selection mask for quantifying the fluorescence intensity in the green and red channels. Oxidation of BODIPY C11 581/591 was calculated as the ratio of the green (fluorescence emission of the oxidized probe)/ red fluorescence mean intensity (fluorescence emission of reduced probe) within the cell outlines.

### Lipidomic analysis

Non-targeted lipidomic analysis of major lipids was performed by reversed-phase ultra-high-performance liquid chromatography (RP-UHPLC) coupled to electrospray ionization time-of-flight mass spectrometry (ESI-TOFMS). Prior to lipid extraction, a mixture of lipid internal standards (**Table 2**) was added to the samples for semi-quantification of reported lipid molecular species. Lipid extraction was performed according to a method adapted from^62^. 500.000 cells were homogenized in 500 μL of 50 mM phosphate buffer (pH 7.4) containing 100 μM deferoxamine mesylate. This homogenate was mixed with 400 μL of ice-cold methanol, containing 100 μM of butylated hydroxytoluene (BHT), and 100 μL of internal standards (10 µg/mL). 2 mL of chloroform: ethyl acetate (4:1) were added to the mixture, followed by vortexing during 30 s. After centrifugation at 1,500 x g for 2 min at 4°C, the lower phase containing total lipid extracts (TLE) was transferred to a new tube and dried under N_2_ gas^63^. Dried TLE were dissolved in 100 µL of isopropanol and the UHPLC injection volume was set at 2 µL. The separation conditions of mass spectrometry analysis were performed as previously described. The MS/MS data were analysed with PeakView®, and lipid molecular species were identified by an in-house manufactured Excel-based macro. Lipids were named according to the LIPID MAPS® Structure Database (LMSD) guidelines^64,65^. The lipid quantification was performed with MultiQuant®, in which peak areas of precursor ions were normalized to those of the internal standards. Final data were expressed as mass of lipid species per mass of total proteins, determined by BCA Protein Assay Kit (Thermo) as manufacturer instructions. Lipids were annotated according to their lipid subclass. Individual lipids were also grouped as the total number of double bonds in: saturated (no double bounds); monounsaturated (presence of one double bound) or polyunsaturated (presence of more than one double bound).

**Table 2:** Lipid internal standards (from Avanti Polar Lipids Inc., Alabaster, Alabama, USA)

| *Internal standards* | *Lipid* | *Concentration*<br>*(ng/μL)* |
| --- | --- | --- |
| cholest-5-en-3β-yl (decanoate) | CE 10:0 | 10 |
| N-decanoyl-D-erythro-sphingosine | Cer d18:1/10:0 | 10 |
| N-heptadecanoyl-D-erythro-sphingosine | Cer d18:1/17:0 | 10 |
| 1',3'-bis[1,2-dimyristoyl-sn-glycero-3-phospho]-glycerol | CL 14:0 x 4 | 10 |
| 1-heptadecanoyl-2-hydroxy-sn-glycero-3-phosphocholine | LPC 17:0 | 10 |
| 1-(10Z-heptadecenoyl)-sn-glycero-3-phosphoethanolamine | LPE 17:1 | 10 |
| 1,2-diheptadecanoyl-sn-glycero-3-phosphate | PA 17:0/17:0 | 10 |
| 1,2-dimyristoyl-sn-glycero-3-phosphocholine | PC 14:0/14:0 | 10 |
| 1,2-diheptadecanoyl-sn-glycero-3-phosphocholine | PC 17:0/17:0 | 10 |
| 1,2-dimyristoyl-sn-glycero-3-phosphoethanolamine | PE 14:0/14:0 | 10 |
| 1,2-diheptadecanoyl-sn-glycero-3-phosphoethanolamine | PE 17:0/17:0 | 10 |
| 1,2-diheptadecanoyl-sn-glycero-3-phospho-(1'-rac-glycerol) | PG 17:0/17:0 | 10 |
| 1,2-diheptadecanoyl-sn-glycero-3-phospho-L-serine | PS 17:0/17:0 | 10 |
| N-heptadecanoyl-D-erythro-sphingosylphosphorylcholine | SG d18:1/17:0 | 10 |
| 1,2,3-tritetradecanoyl-sn-glycerol | TG 14:0 x 3 | 10 |
| 1,2,3-triheptadecanoyl-sn-glycerol | TG 17:0 x 3 | 10 |

### Labile iron pool (LIP) measurement

LIP was given as sum of the concentrations of iron ([Fe]) and calcein-bound Fe ([CA-Fe]), normalized to the total intracellular calcein ([CA]t), whereby LIPN = LIP/[CA]t. We followed the rationale for fluorescence determination of LIP developed by^66^ with minor modifications. Cells, at a density of 1-2 × 10^6^ cells/mL, were incubated with 0.25 µM of calcein acetoxymethyl ester (CA–AM) for 5 min at 37°C in bicarbonate-free and serum-free growth medium containing 1 mg/mL BSA and 20 mM Hepes (Sigma-Aldrich), pH 7.3. After incubation, cells were washed of excess CA-AM with medium without CA-AM 2 times, and resuspended in warm HBS (Hepes 20 mM, NaCl 150 mM, pH 7.3). The number of cells (Nc) was measured by counting in a hemocytometer. The cell suspension was transferred to a 24-well microplate and fluorescence of calcein (CA)-loaded cells (F) was monitored at an excitation of 488 nm and emission of 517 nm using a microplate reader (Infinite M200 plate reader-Tecan, Männedorf, Switzerland), with gently orbital shaking before each measurement (2 seconds, 4 mm amplitude). After signal stabilization (2-5 minutes) and reaching a given fluorescence intensity (F), the iron chelator SIH (Salicylaldehyde Isonicotinoyl Hydrazine, 100 µM final) was added, causing a rise in fluorescence signal (Fc). The rise in fluorescence elicited by SIH was given as the fractional change (ΔF), using a normalized fluorescence scale FN = F/Fc. Next, the CA concentration in the cell suspension ([CA]susp) was determined from a calibration curve obtained by adding CA standards (in 1 nM steps) to the CA-loaded cells suspension supplemented with SIH. The [CA]t was calculated from the relationship [CA]t = [CA]susp/Nc. The [CA-Fe] was obtained from the relationship [CA-Fe] = ΔF∗[CA]t. [Fe] was calculated from CA-Fe dissociation constant: *Kd* = [CA]t∗[Fe]/[CA-Fe]), using the experimental values of [CA-Fe] and [CA]t and the *Kd* in cells value of (0.22) obtained in the original paper66. CA, CA-AM and SIH were generous gifts from Dr. Breno Pannia Espósito, Chemistry Institute of the University of São Paulo, Brazil.

### Statistical analysis

All results were analysed for Gaussian distribution and passed the normality test. The statistical differences between group means were tested by One-way ANOVA followed by Tukey’s post-test for multiple comparisons or by Two-way ANOVA followed by Bonferroni’s post-test for multiple comparisons. For PCA in lipidomic studies, statistical analysis was performed with MetaboAnalyst website. A value of p<0.05 was considered as statistically significant in all analysis. All data presented in this manuscript are available upon request to the authors.

## Acknowledgements

This research was funded by the Brazilian agencies FAPESP (grants 2019/09517-2, 2019/05026-4, 2017/18922-2, 2017/03618-6, 2017/13804-1, 2016/04676-7, 2015/02654-3, 2013/07937-8), CAPES and CNPq. We are very grateful to prof. Dr. Breno Pannia Espósito, from IQ-USP, for the kind help and practical suggestions for LIP measurement. We also kindly acknowledge the support of the lab technicians Marcelo S. Nunes and Sandra R. Souza, and the members of LFPI laboratory, from IQ-USP, specially Dr. Helena C. Junqueira and Dr. Felipe G. Ravagnani.

## Supplementary Figures

**Supplementary Figure 1:**
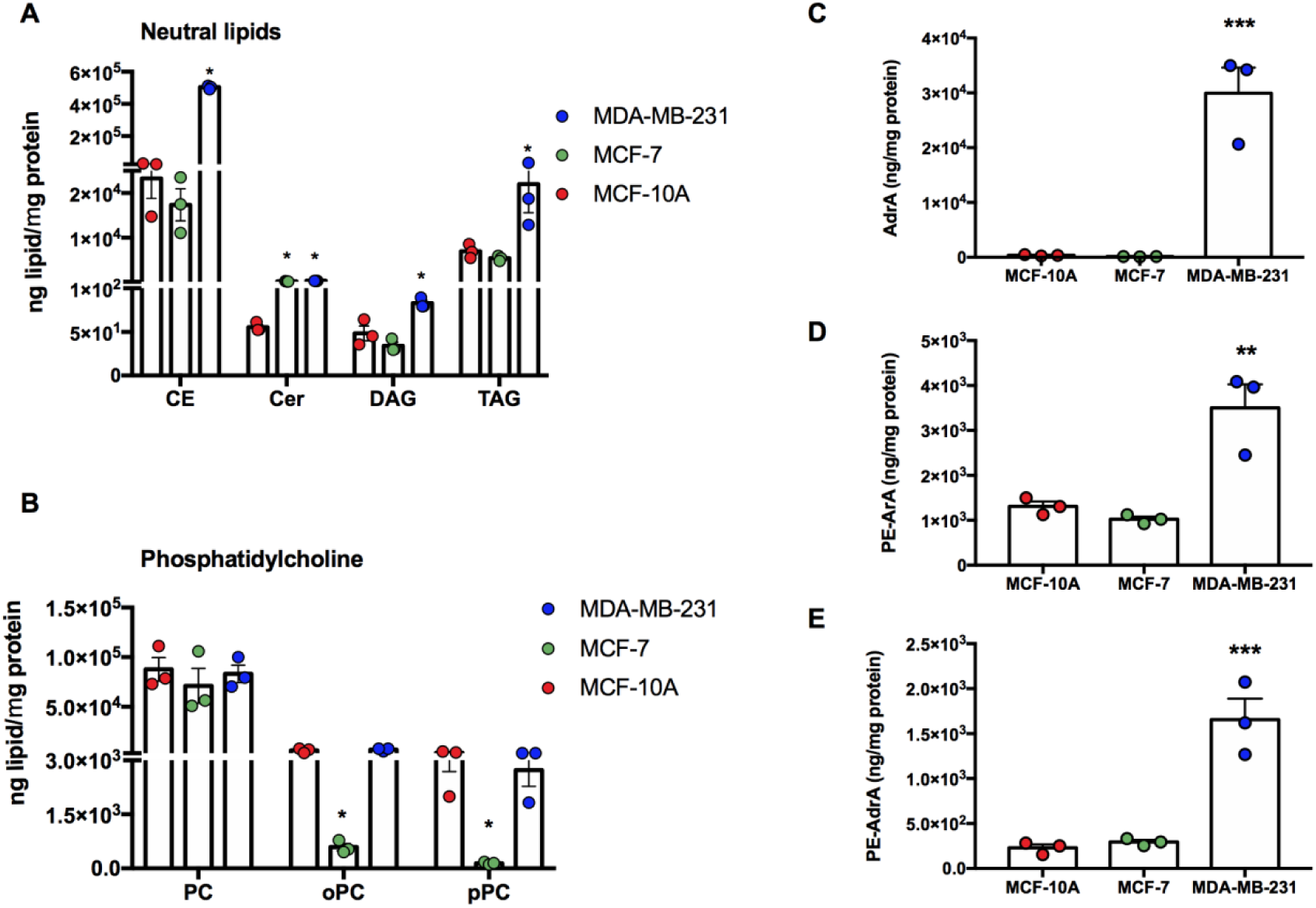
Basal lipid composition of breast cells. Lipids were extracted from the three cell types. (A) Neutral lipid abundance: Cholesteryl Esters (CE), Ceramides (Cer), Diacylglycerides (DG) and Triacylglycerides (TG). (B) Abundance of phosphatidylcholine (PC), plasmanyl (o)- and plasmenyl (p)-PCs (oPC and pPC respectively). (C) Abundance of Adrenic Acid (AdrA)-containing lipids. (D) Abundance of Arachidonic Acid ArA esterified in phosphatidyletanolamine (PE). (E) Abundance of AdrA esterified in PE. *** p<0.001; ** p<0.005; *p<0.05 *vs* MCF-10A. Results are presented as mean ± S.E.M. n=3 independent experiments. Dot colors representation: MCF-10A in red; MCF-7 in green; MDA-MB-231 in blue.

**Supplementary Figure 2:**
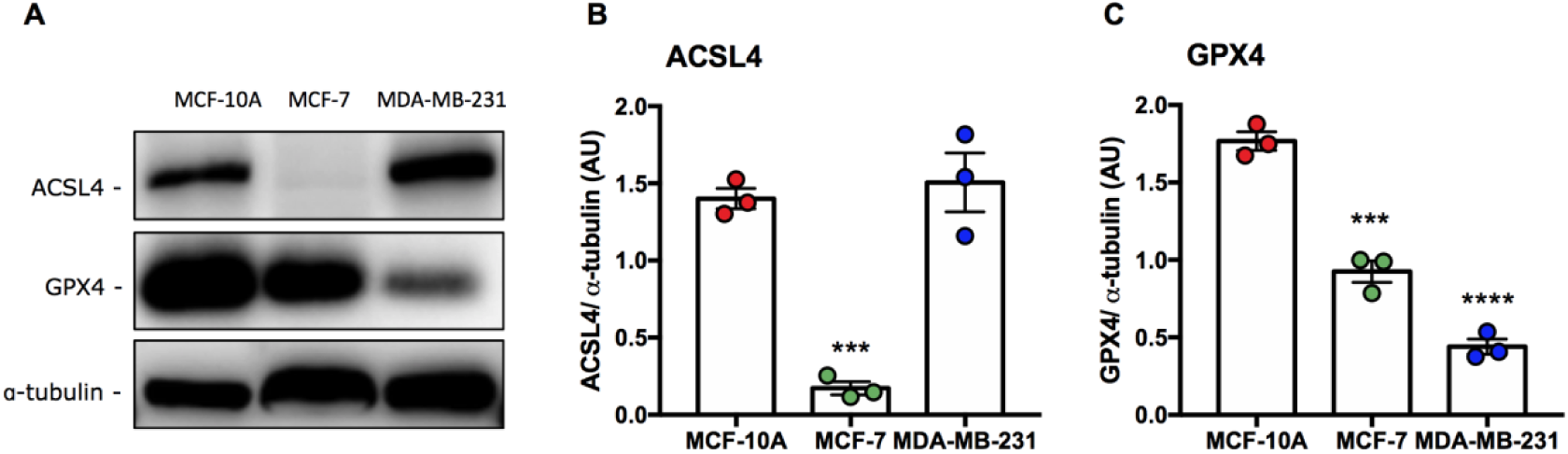
Basal abundance of ferroptosis key players. (A) Representative pictures of Western blots and the corresponding quantification of (B) ACSL4 and (C) GPX4. **** p<0.0001; *** p<0.001 *vs* MCF-10A. Results are presented as mean ± S.E.M. n=3 independent experiments. Dot colors representation: MCF-10A in red; MCF-7 in green; MDA-MB-231 in blue.

**Supplementary Figure 3:**
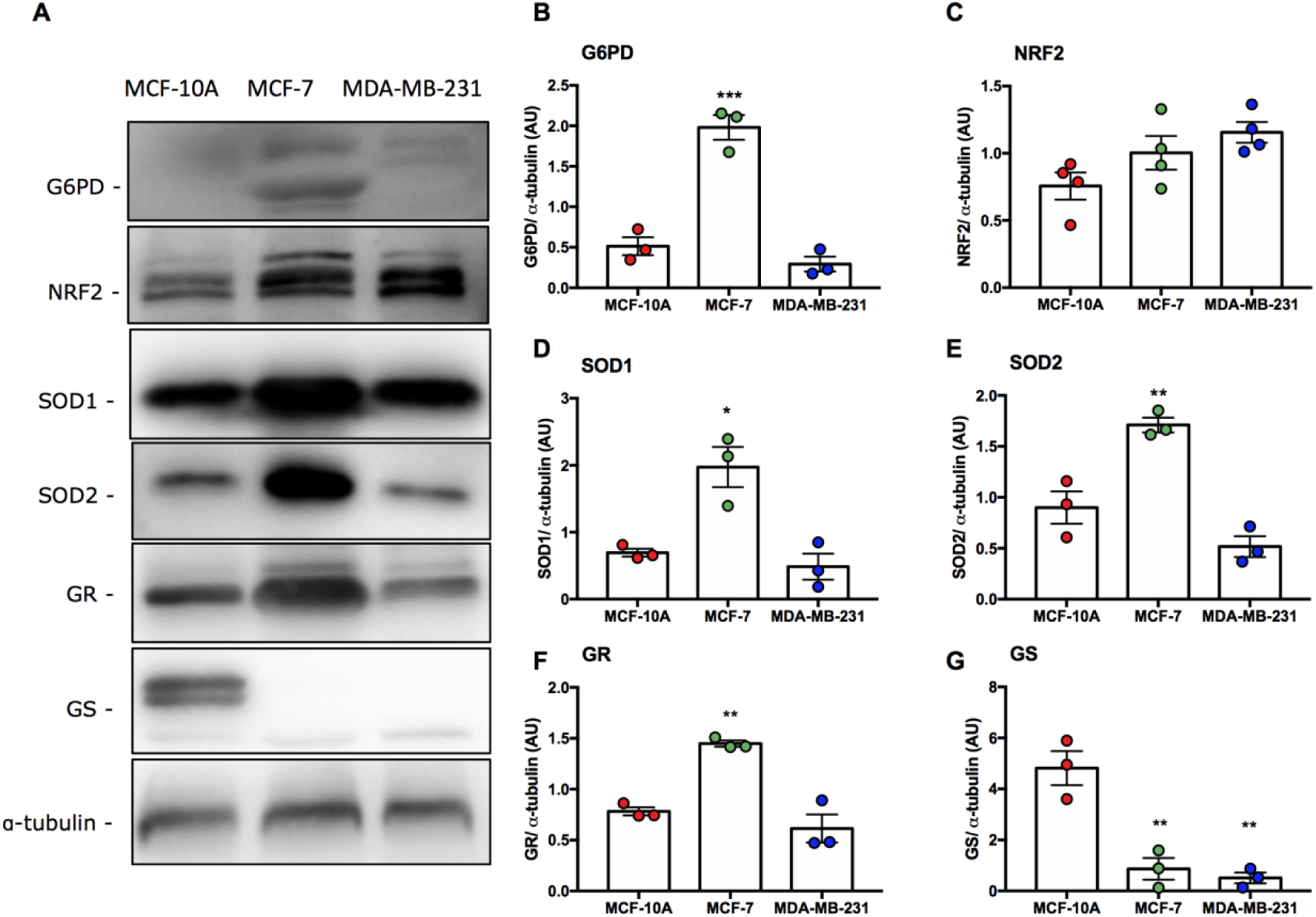
Basal abundance of antioxidant-related proteins. (A) Representative pictures of Western blots and the corresponding quantification of (B) G6PD, (C) NRF2, (D) SOD1, (E) SOD2, (F) GR- glutathione reductase and (G) GS-glutathione synthetase. *** p<0.001 *vs* MCF-10A. Results are presented as mean ± S.E.M. n≥3 independent experiments. Dot colors representation: MCF-10A in red; MCF-7 in green; MDA-MB-231 in blue.

**Supplementary Figure 4:**
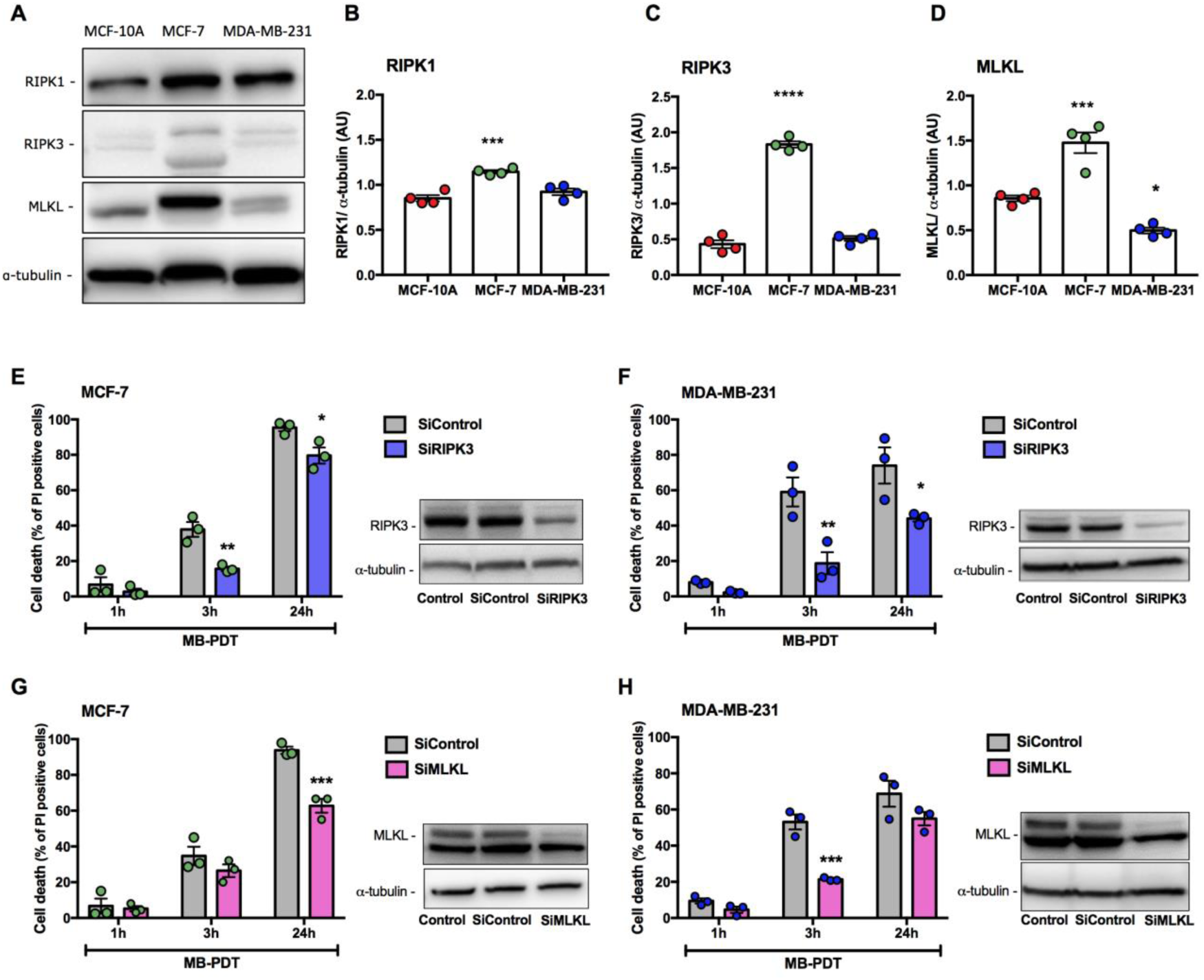
MB-PDT induces necroptosis in breast tumor cells. (A) Representative pictures of Western blots and the corresponding quantification of basal levels of (B) RIPK1, (C) RIPK3, (D) MLKL. (E, F) RIPK3 or (G, H) MLKL was silenced or not in the cells, which were then submitted to MB-PDT. Cells were submitted or not to MB-PDT and cell death was analysed after 1, 3 or 24h of cell treatment. Right panels: representative Western blot analysis of RIPK3 and MLKL protein levels. **** p<0.0001; *** p<0.001; ** p<0.005; *p<0.05 *vs* MCF-10A. Results are presented as mean ± S.E.M. n≥3 independent experiments.

