## Supplementary Material for "Distinct photooxidation-induced cell death pathways lead to selective killing of human breast cancer cells"

### Supplementary Figures

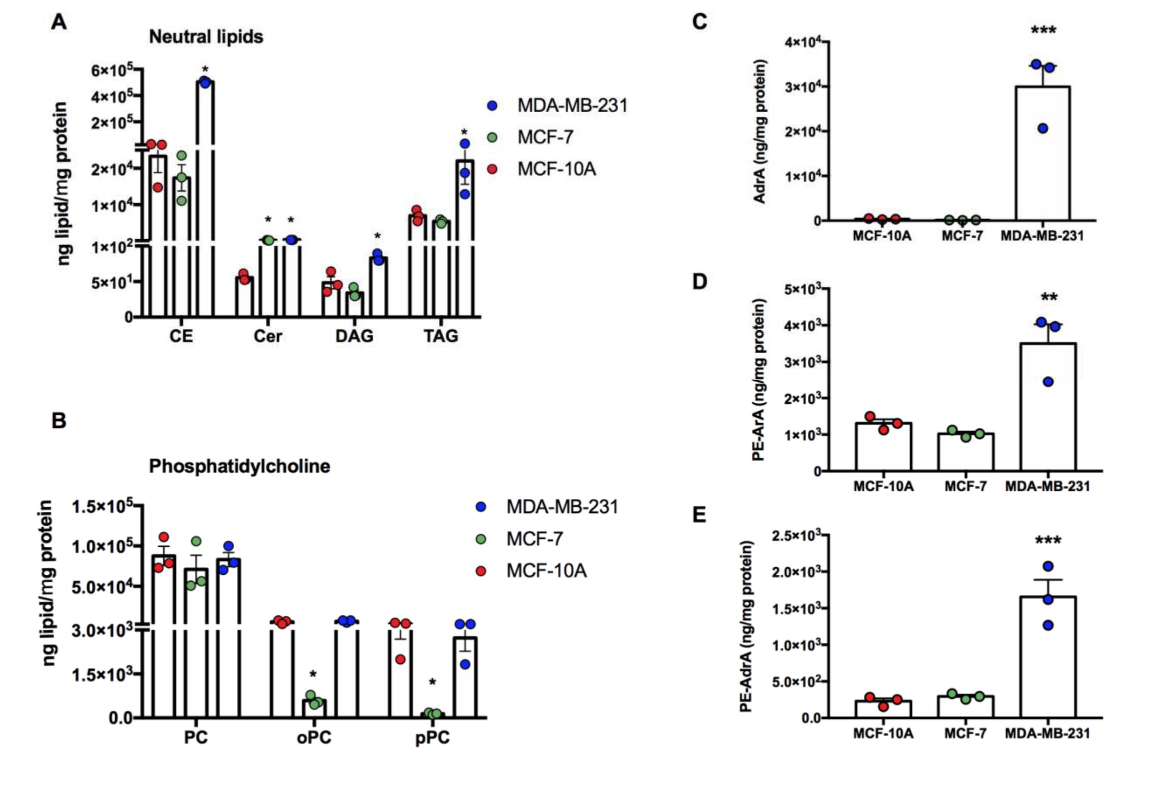


**Supplementary Figure 1: Basal lipid composition of breast cells.** Lipids were extracted from the three cell types. (A) Neutral lipid abundance: Cholesteryl Esters (CE), Ceramides (Cer), Diacylglycerides (DG) and Triacylglycerides (TG). (B) Abundance of phosphatidylcholine (PC), plasmanyl (o)- and plasmenyl (p)-PCs (oPC and pPC respectively). (C) Abundance of Adrenic Acid (AdrA)-containing lipids. (D) Abundance of Arachidonic Acid ArA esterified in phosphatidyletanolamine (PE). (E) Abundance of AdrA esterified in PE. *** $p<0.001$; ** $p<0.005$; *$p<0.05$ *vs* MCF-10A. Results are presented as mean ± S.E.M. n=3 independent experiments. Dot colors representation: MCF-10A in red; MCF-7 in green; MDA-MB-231 in blue.

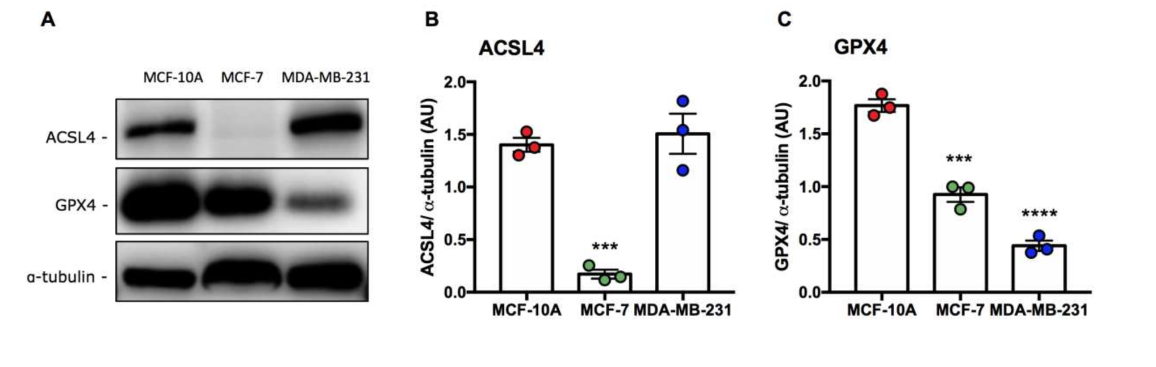


**Supplementary Figure 2: Basal abundance of ferroptosis key players.** (A) Representative pictures of Western blots and the corresponding quantification of (B) ACSL4 and (C) GPX4. **** $p<0.0001$; *** $p<0.001$ *vs* MCF-10A. Results are presented as mean ± S.E.M. n=3 independent experiments. Dot colors representation: MCF-10A in red; MCF-7 in green; MDA-MB-231 in blue.

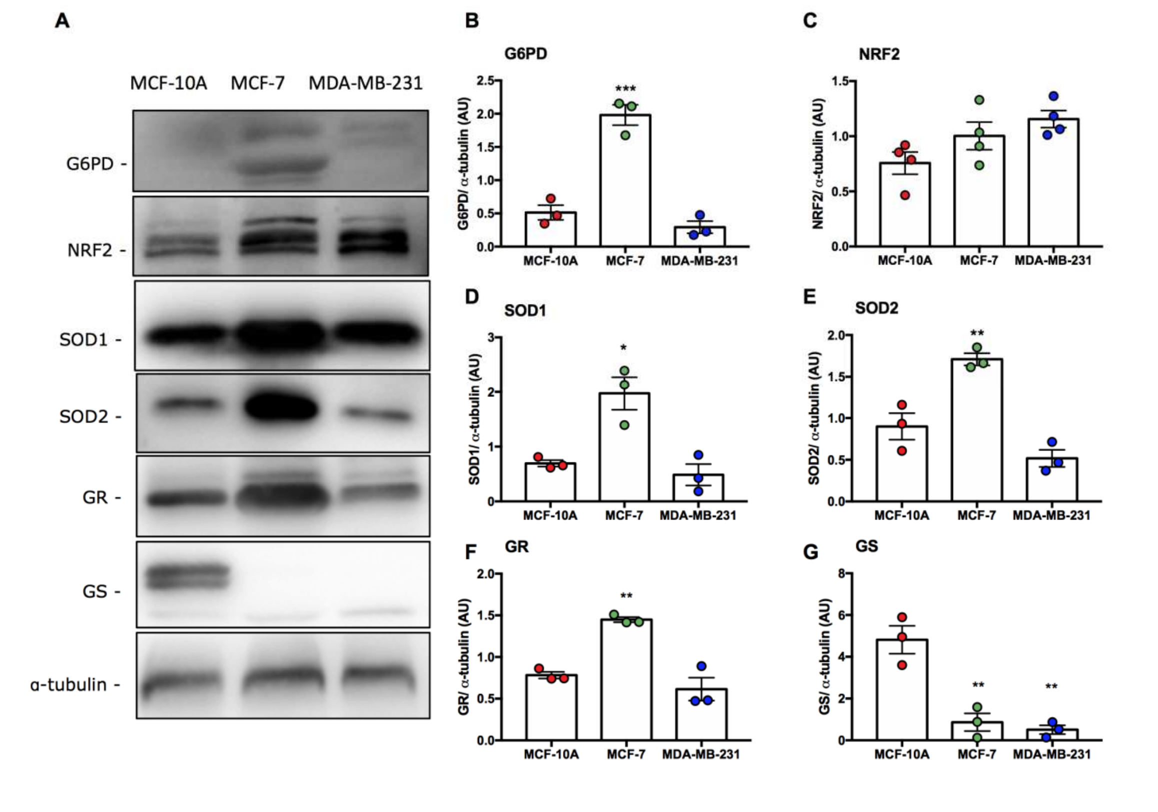


**Supplementary Figure 3: Basal abundance of antioxidant-related proteins.** (A) Representative pictures of Western blots and the corresponding quantification of (B) G6PD, (C) NRF2, (D) SOD1, (E) SOD2, (F) GR-glutathione reductase and (G) GS-glutathione synthetase. *** $p<0.001$ *vs* MCF-10A. Results are presented as mean ± S.E.M. $n\geq3$ independent experiments. Dot colors representation: MCF-10A in red; MCF-7 in green; MDA-MB-231 in blue.

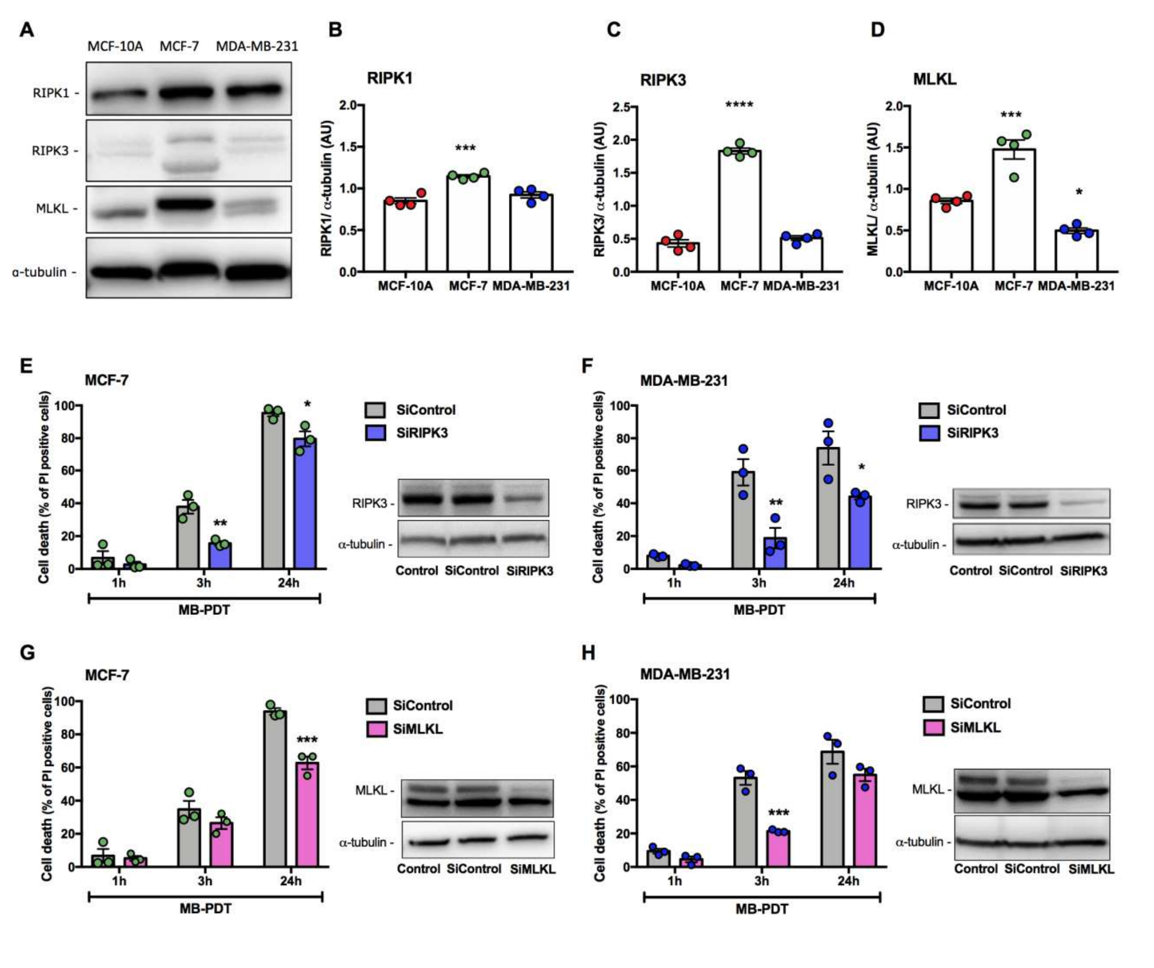


Supplementary Figure 4: MB-PDT induces necroptosis in breast tumor cells. (A) Representative pictures of Western blots and the corresponding quantification of basal levels of (B) RIPK1, (C) RIPK3, (D) MLKL. (E, F) RIPK3 or (G, H) MLKL was silenced or not in the cells, which were then submitted to MB-PDT. Cells were submitted or not to MB-PDT and cell death was analysed after 1, 3 or 24h of cell treatment. Right panels: representative Western blot analysis of RIPK3 and MLKL protein levels. **** p<0.0001; *** p<0.001; ** p<0.005; *p<0.05 *vs* MCF-10A. Results are presented as mean ± S.E.M. n≥3 independent experiments.
